# *In vivo* CRISPR screening identifies *NF1/RASA1/TP53* co-mutations and downstream MEK signaling as a common key mechanism of sinonasal tumorigenesis

**DOI:** 10.1101/2025.05.19.654661

**Authors:** Kenny Vu, Sreenivasulu Gunti, Ramya Viswanathan, Anjali Nandal, Riley Larkin, Sungwoo Cho, Jonathan Zou, Shivani Ramolia, Austin T.K. Hoke, Siani M. Barbosa, Gary L. Gallia, Lisa M. Rooper, Charalampos S. Floudas, Hui Cheng, Christine N. Miller, Mary R. Guest, Marco Notaro, Arati Raziuddin, Zhonghe Sun, Xiaolin Wu, Farhoud Faraji, Matt Lechner, Federico Comoglio, Elijah F. Edmondson, Raj Chari, Nyall R. London

## Abstract

Genomic alterations driving tumorigenesis in sinonasal malignancies remain largely unexplored. Here, we perform an *in vivo* loss-of-function screen using a pooled custom single-guide library delivered to the sinonasal cavity by adeno-associated virus vector to identify cancer driver genes across diverse sinonasal malignancies. This approach yielded sinonasal malignancies with diverse histologies, including sinonasal squamous cell carcinoma, adenocarcinoma, poorly differentiated sinonasal carcinoma, and sinonasal neuroendocrine tumors characteristic of olfactory neuroblastoma. Surprisingly, rather than observing distinct sgRNA profiles across sinonasal tumor subtypes, common recurrent mutations were identified in *Nf1* (79%), *Rasa1* (74%), and *Trp53* (68%) across malignancies with distinct histologies. Utilizing an orthogonal approach, we confirmed that *Nf1/Trp53* were required for sinonasal tumorigenesis. Given that loss-of-function in *NF1* and *RASA1* may lead to increased Ras activity and downstream MEK signaling, we tested small molecule targeting of the RAS-MAPK pathway in sinonasal malignancies. Indeed, both tumor cell lines derived from our loss-of-function approach as well as from human sinonasal malignancies displayed significant sensitivity to MEK inhibition in standard *in vitro* culture and organoid models. These findings demonstrate that loss of NF1 and RASA1-mediated Ras-GAP activity leads to Ras activation and downstream MEK signaling which is a potential common target throughout major sinonasal tumor subtypes.

## Main

The diversity of tumor histopathology in the sinonasal cavity is amongst the highest in the human body^1,2^. This heterogeneity is in part due to a distinctive tissue composition including a unique olfactory neuroepithelium, respiratory mucosa, minor salivary glands, and others which directly interface with the external airborne environment. Sinonasal malignancies are rare and comprise <1% of all cancer cases^2^. These most commonly include sinonasal squamous cell carcinoma (SNSCC), sinonasal adenocarcinoma, neuroendocrine tumors such as olfactory neuroblastoma (ONB), and poorly differentiated carcinomas such as sinonasal undifferentiated carcinoma (SNUC)^3,4^. Treatment varies depending on the tumor type but typically includes multi-modality therapy such as surgery, radiation, and/or chemotherapy to achieve optimal outcomes^2,5,6^. However, these treatment approaches may cause considerable morbidity due to the close proximity to critical neurovascular structures such as cranial nerves, the eyes, and brain^7,8^.

Due to the rare nature of sinonasal malignancies, mechanisms of tumorigenesis are poorly understood, and genomics studies have been limited by small patient cohorts. *TP53* is the most frequently mutated gene amongst many sinonasal tumor types, but the identification of common molecular mechanisms governing sinonasal oncogenesis is lacking^9–13^. Indeed, *in vivo* screening approaches to identify genes driving sinonasal tumorigenesis have not been previously reported. We hypothesized that *in vivo* CRISPR screening would elucidate mechanisms of tumorigenesis across sinonasal cancer types and identify potential therapeutic targets.

### *In vivo* CRISPR screening generates a variety of histologic sinonasal tumor subtypes

We generated a pooled single-guide RNA (sgRNA) library including 167 genes selected from genes implicated in sinonasal and head and neck cancer tumorigenesis, and prior *in vivo* CRISPR screening approaches in other organs (**Supplementary Table 1**)^11,14,15^. Given the frequency of *TP53* mutations amongst sinonasal tumor types^9–12^, a *TP53* sgRNA was included in the backbone of the library construct. Next, we utilized a variety of adeno-associated virus (AAV) serotypes to determine which provides the optimal viral transduction of a variety of cell types in the murine sinonasal cavity. Through instillation of a variety of AAV-Cre serotypes in the sinonasal cavity of Rosa26-LacZ reporter (R26R) mice, AAV5 was identified as the ideal serotype to achieve the goal of broad transduction across both the olfactory and respiratory epithelium (**Supplementary Figure 1A-B**). The pooled sgRNA library was therefore packaged in AAV5 and instilled thrice in the sinonasal cavity of Cas9 transgenic (H11*^Cas9^*) mice.

AAV5-*TP53*-sgRNA library treated H11*^Cas9^* mice were monitored with microCT and developed sinonasal tumors approximately 3-5 months after instillation (**Figure 1; Supplementary Figure 2**). As tumors enlarged in size, destruction of turbinate structures was noted followed by extranasal invasion and growth (**Figure 1; Supplementary Figure 2**). Of the 33 H11*^Cas9^* mice treated with the sgRNA library 28 mice formed sinonasal tumors, while the remaining 5 mice succumbed to other causes prior to reaching 6 months post-AAV5 exposure. This was in contrast to control H11*^Cas9^* mice treated with the AAV5 viral backbone alone (AAV5-*TP53*-null) in which none of the 27 control mice formed sinonasal tumors up to 16 months after treatment (**Supplementary Figure 3**). Next, we sought to determine the histologic subtype of each sinonasal tumor. Tumor tissue was collected and when sufficiently available, underwent H&E staining as well as immunohistochemistry analysis for a variety of known markers of sinonasal malignancies including p63 and cytokeratin and included known neuroendocrine markers of sinonasal malignancies including synaptophysin (SYP), neuron-specific enolase (NSE), and chromogranin. Tissues were assessed by a veterinary pathologist and a variety of sinonasal tumor types including the major sinonasal subtypes were identified namely SNSCC, adenocarcinoma, poorly differentiated sinonasal tumors similar to sinonasal undifferentiated carcinoma (SNUC), and sinonasal neuroendocrine tumors characteristic of olfactory neuroblastoma (ONB) (**Figure 2, Supplementary Figure 4**). Thus, the major sinonasal tumor types could be generated using a pooled AAV5-*TP53*-sgRNA screening approach (**Table 1**).

**Figure 1.**
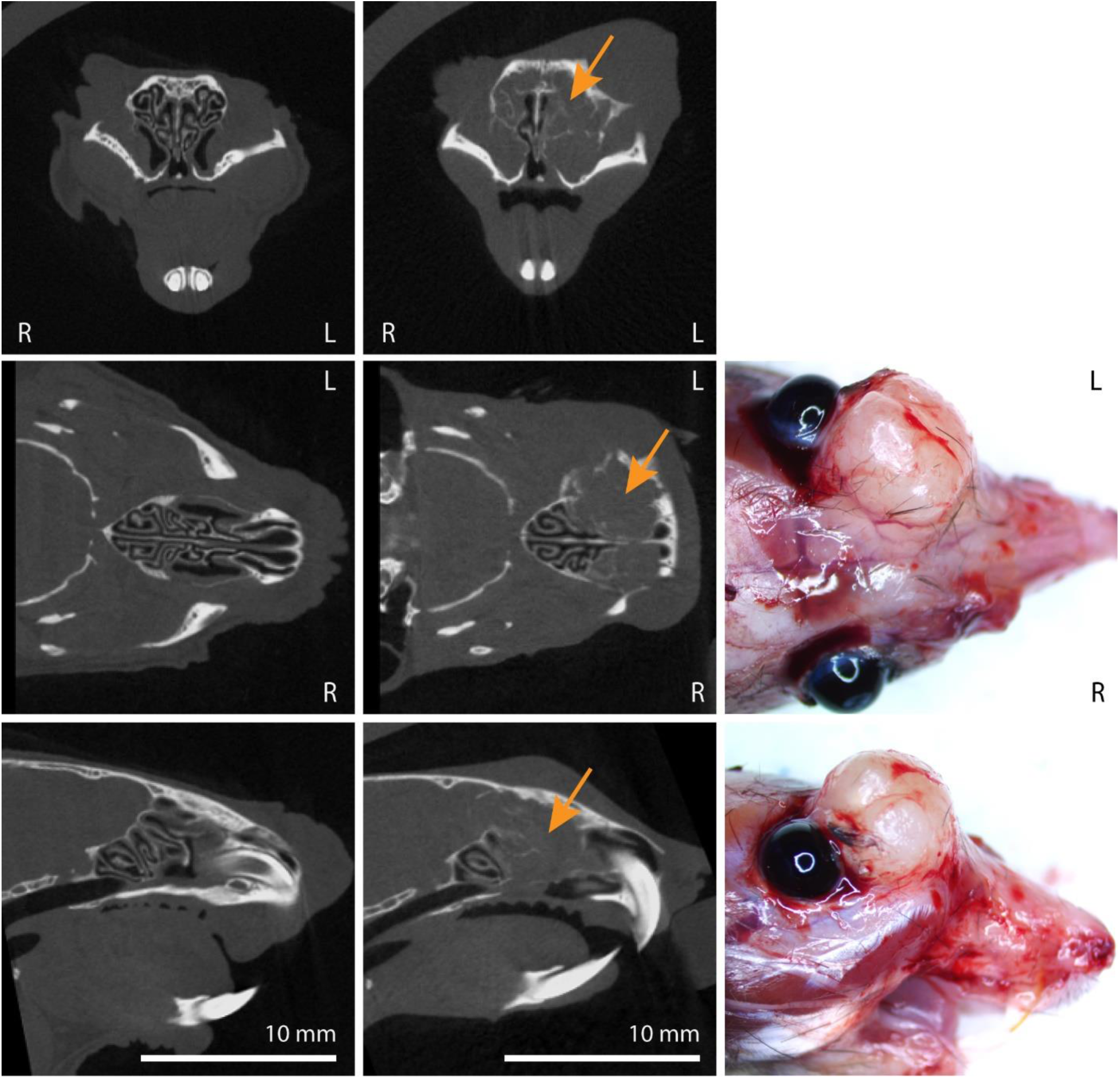
Sinonasal tumor formation after AAV5-*TP53*-sgRNA library instillation in H11*^Cas9^* mice. (Left column) AAV5-*TP53*-null control treated H11*^Cas9^* mice. MicroCT images in the coronal (top), axial (middle), and sagittal (bottom) planes demonstrate normal turbinate structure. **(Middle column)** AAV5-*TP53*-sgRNA library treated H11*^Cas9^* mice. MicroCT images in the coronal (top), axial (middle), and sagittal (bottom) planes demonstrate representative tumor formation (orange arrows) and destruction of sinonasal turbinates. **(Right column)** AAV5-*TP53*-sgRNA library treated H11*^Cas9^* mice. Demonstration of representative extranasal invasion and growth in the axial (middle) and sagittal (bottom) planes.

**Figure 2.**
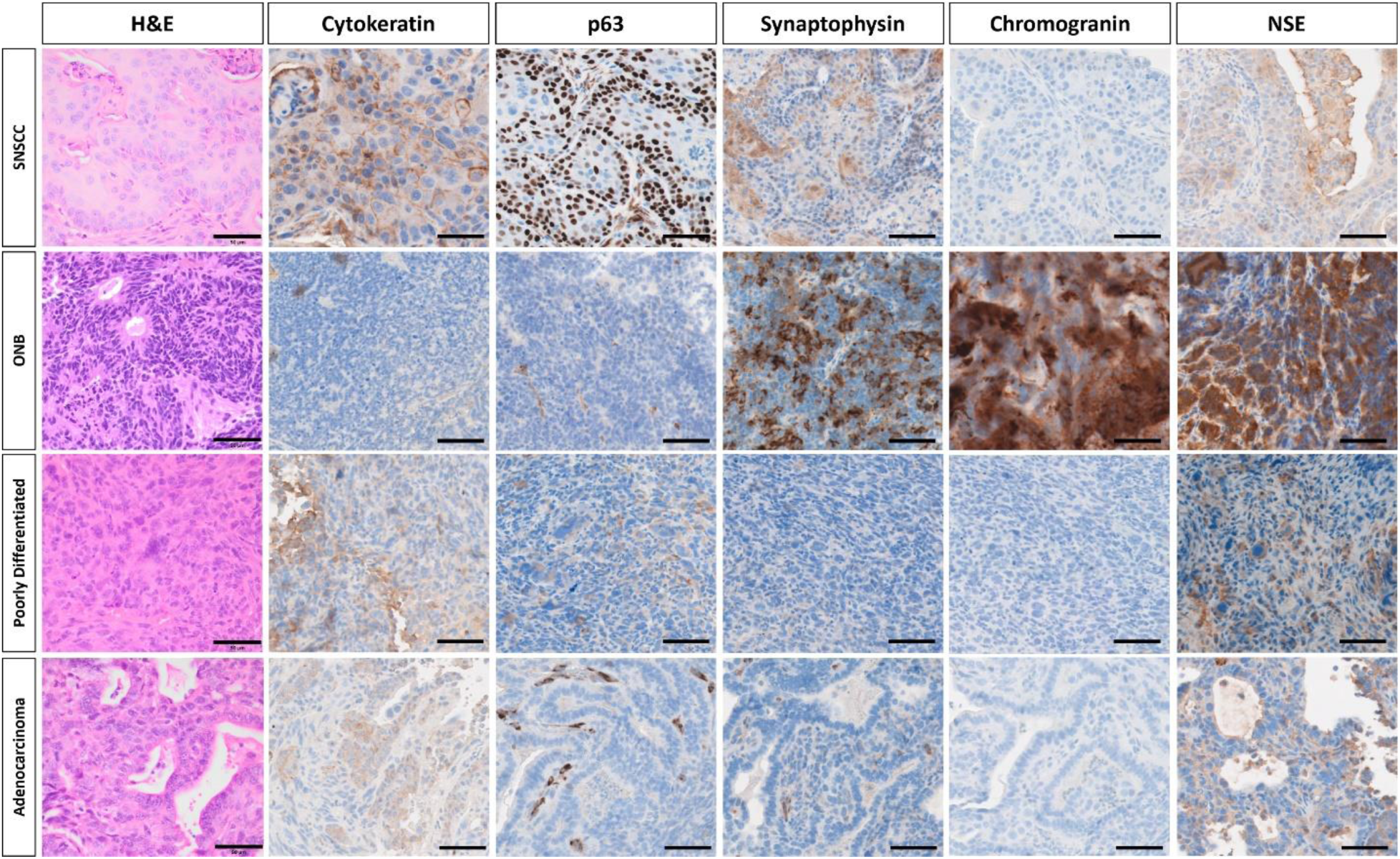
Four distinct types of sinonasal tumors were identified on histologic characterization of murine sinonasal tumors. (Top row) SNSCC tumors were positive for cytokeratin and p63 and demonstrated haphazardly arranged trabeculae of neoplastic cells with squamous differentiation, including large cells with eosinophilic cytoplasm, prominent intracellular bridges, and keratohyalin granules with rare dyskeratosis and keratin pearl formation. **(Second row)** ONB were positive for synaptophysin, chromogranin, and NSE and demonstrated densely cellular tumors composed of nests of cells supported by a delicate fibrovascular stroma. Cells were small, round to columnar, and have scant cytoplasm containing oval, basally-located nuclei. Neoplastic cells form rosettes, characterized by taller columnar cells surrounding a central lumen containing eosinophilic fibrillar debris. **(Third row)** Poorly differentiated tumors demonstrated sheets of poorly differentiated neoplastic cells with minimal stroma and high mitotic activity. Neoplastic cells are pleomorphic and karyomegalic cells containing nuclear profiles 3-5x larger than neighboring tumor cells are present. Sarcomatoid morphology is multifocally present in some cases, characterized by spindloid cells forming interlacing bundles. **(Fourth row)** Sinonasal adenocarcinoma tumors demonstrated mild cytokeratin expression and tumors were composed of malignant epithelial cells forming irregular glands and acini composed of cuboidal cells. Representative images of each tumor types are shown. Scalebar equals 50µm. SNSCC – sinonasal squamous cell carcinoma, ONB – olfactory neuroblastoma, H&E – hematoxylin & eosin, NSE – neuron-specific enolase.

**Table 1.** Histological characterization of murine sinonasal tumors. . * Indicates NF1/P53^flox/flox^ background. CK (cytokeratin), SYP (synaptophysin), NSE (neuron-specific enolase), CgA (chromogranin A).

| **Tumor** | **Histology Type** | **CK** | **p63** | **SYP** | **CgA** | **NSE** |
| --- | --- | --- | --- | --- | --- | --- |
| **1** | Squamous cell carcinoma | +++ | +++ | - | - | + |
| **2** | Olfactory neuroblastoma | - | - | ++ | - | ++ |
| **3\*** | Olfactory neuroblastoma | - | - | +++ | ++ | + |
| **5** | Poorly differentiated carcinoma/SNUC | ++ | - | - | - | - |
| **7** | Squamous cell carcinoma | ++ | +++ | - | - | ++ |
| **8** | Squamous cell carcinoma | ++ | ++ | + | - | + |
| **10** | Poorly differentiated carcinoma/SNUC | + | - | - | - | - |
| **14** | Adenocarcinoma | ++ | + | - | - | - |
| **15** | Adenocarcinoma | ++ | + | - | - | ++ |
| **45** | Adenocarcinoma | ++ | +++ | - | - | - |
| **48** | Squamous cell carcinoma | ++ | +++ | - | - | - |
| **49** | Adenocarcinoma | ++ | - | - | - | - |
| **50** | Poorly differentiated carcinoma/SNUC | ++ | - | - | - | + |
| **52\*** | Olfactory neuroblastoma | - | - | ++ | + | ++ |
| **53** | Poorly differentiated carcinoma/SNUC | ++ | - | + | - | + |
| **57** | Adenocarcinoma | ++ | - | - | - | ++ |

### Frequent *NF1/RASA1/TP53* co-mutations identified amongst sinonasal tumor subtypes

Next, we sought to identify which sgRNAs were responsible for tumor development in each sinonasal tumor. Sufficient tumor tissue was available for 19 sinonasal tumors derived from H11*^Cas9^* mice. A custom capture library was applied followed by next-generation sequencing to identify the inciting sgRNAs for each tumor. Surprisingly, rather than unique, distinct sgRNA mutational profiles across sinonasal tumor subtypes, common recurrent mutations were identified in *NF1* (79%), *RASA1* (74%), and *TP53* (68%) (**Figure 3**). Interestingly, both *NF1* and *RASA1* sgRNA co-occurred in 13/19 (68%) tumors, many of which also had *TP53* mutations. Of the remaining 6 tumors, all had *TP53* sgRNA and 3/6 (50%) had either an *NF1* or *RASA1* sgRNA target. Collectively these data demonstrate that *NF1*/*RASA1*/*TP53* co-mutations are common key mechanisms of sinonasal tumorigenesis across the major sinonasal tumor types in mice.

**Figure 3.**
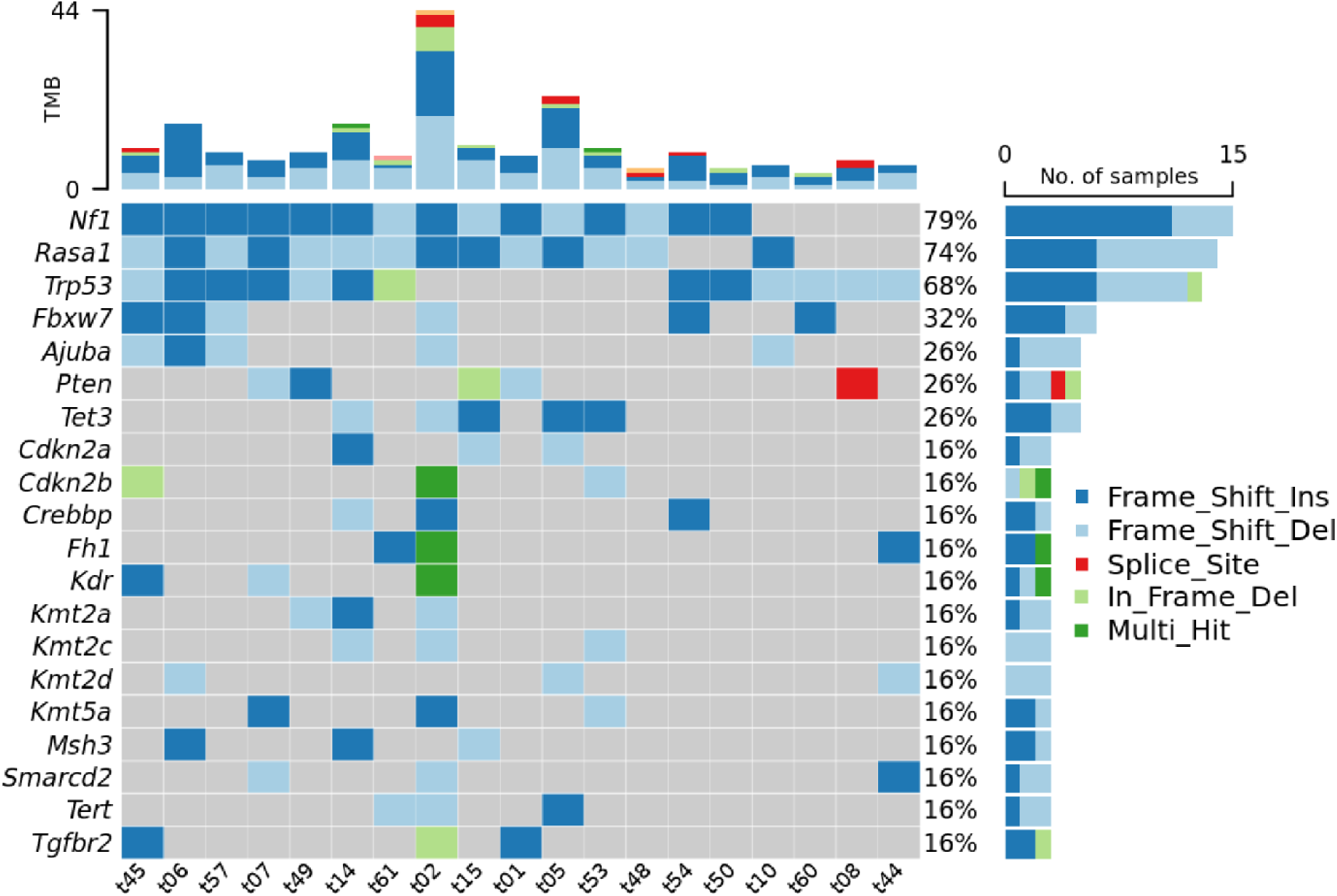
Frequent *NF1/RASA1/TP53* co-mutations identified amongst the four sinonasal tumor types. Murine sinonasal tissue underwent custom capture library directed next-generation sequencing to identify inciting sgRNA for each tumor. *NF1/RASA1/TP53* co-mutations were frequently observed amongst SNSCC, ONB, poorly differentiated carcinoma, and adenocarcinoma tumor types. TMB – tumor mutational burden.

We next evaluated genes with a lower frequency of sgRNA perturbation. Genes involved in DNA or histone methylation were affected in 13 of 19 (68%) of sinonasal tumors and included Tet methylcytosine dioxygenase 3 (*TET3*), DNA (cytosine-5)-methyltransferase 3A (*DNMT3A*), lysine N-methyltransferase (KMT) family members (including *KMT2A*, *KMT2B*, *KMT2C*, *KMT2D*, and *KMT5A*), and lysine demethylase (KDM) family members (*KDM5A*, *KDM5C*, and *KDM6A*) (**Figure 3, Supplementary Table 2**). Similarly, SWItch/Sucrose Non-Fermentable (SWI/SNF) chromatin remodeling complex family members were affected in 13 of 19 (68%) sinonasal tumors including *ARID1A*, *ARID1B*, *SMARCA4*, *SMARCB1*, *SMARCC1*, *SMARCC2*, *SMARCD1*, *SMARCD2*, and *SMARCD3* (**Figure 3, Supplementary Table 2**). Thus, perturbation of DNA or histone methylation and chromatin remodeling pathways may also play a role in sinonasal tumorigenesis.

### Sinonasal tumor development in *NF1/TP53^flox/flox^* mice

To support the *in vivo* CRISPR screen results through a second approach, we next tested whether instillation of viral Cre recombinase vectors could incite sinonasal tumors in *NF1/TP53^flox/flox^* mice. These two genes were selected as *NF1* sgRNA was the most commonly observed affected gene in the screen and *TP53* is the most commonly reported mutated gene in many human sinonasal tumors^9–12^. *NF1/TP53^flox/flox^* mice exposed to AAV5-Cre recombinase administration developed sinonasal tumors (13 of 20 mice) but these tumors were typically much smaller and were first noted later around 12 months on microCT imaging after viral exposure (**Figure 4; Supplementary Figure 5**). No tumor development was noted in 15 *NF1/TP53^flox/flox^*mice exposed to viral null vectors after up to 16 months (**Supplementary Figure 6**). As these tumors tended to be much smaller, thorough immunohistochemical evaluation was much more difficult but sufficient tissue was obtained from 2 larger tumors and tumor histology ascertained (**Table 1**). These results demonstrate that loss of *NF1* and *TP53* alone are sufficient to incite sinonasal tumor development and suggest that additional mutations such as *RASA1*, genes involved in DNA/histone methylation, and SWI/SNF chromatin remodeling family members aid in tumor growth acceleration.

**Figure 4.**
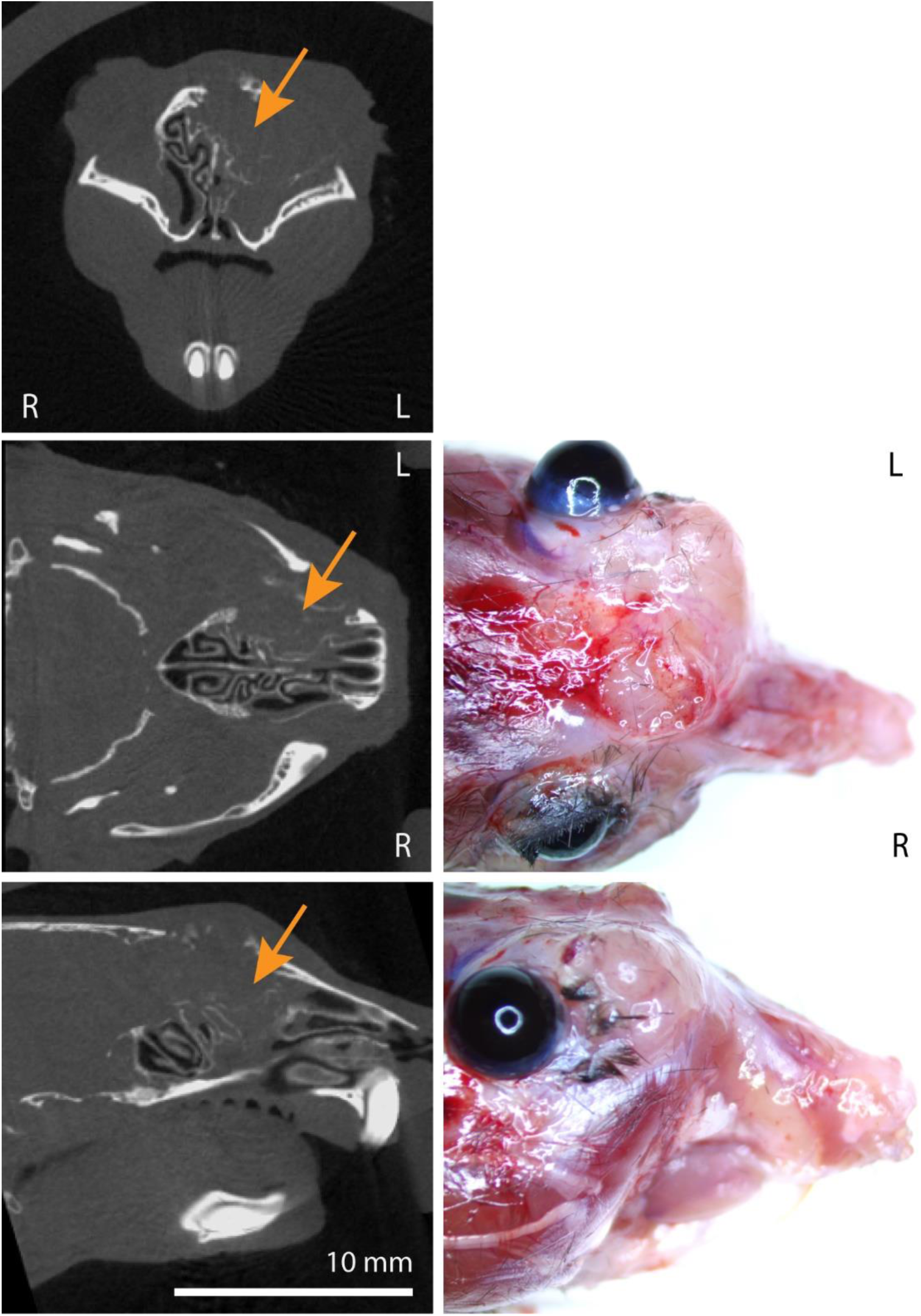
Sinonasal tumor formation after AAV5-Cre instillation in *NF1/TP53^flox/flox^* mice. (Left column) AAV5-Cre treated *NF1/TP53^flox/flox^* mice. MicroCT images in the coronal (top), axial (middle), and sagittal (bottom) planes demonstrate representative tumor formation in a larger tumor (orange arrows) and destruction of sinonasal turbinates. **(Right column)** Demonstration of representative extranasal invasion and growth in the axial (middle) and sagittal (bottom) planes.

### MEK small molecule targeting inhibits sinonasal tumor proliferation

Both NF1 and RASA1 have known Ras-GAP activity, thus loss of these genes may lead to increased Ras function. Ras downstream signaling includes the Raf/MEK/ERK pathway. Thus, we hypothesized that murine sinonasal tumors of varying histologies may be susceptible to MEK inhibition with small molecule MEK inhibitors. First, we generated to our knowledge, the first murine sinonasal tumor cell lines from larger sinonasal tumors for which sufficient tissue was available for each of the 4 major histologic subtypes. These cell lines were subjected to MEK inhibition with Trametinib, Mirdametinib, and Selumetinib. Significant inhibition of tumor cell proliferation *in vitro* was noted with each of these three inhibitors in all 4 murine cell lines (**Figure 5**, *P* < 0.01).

**Figure 5.**
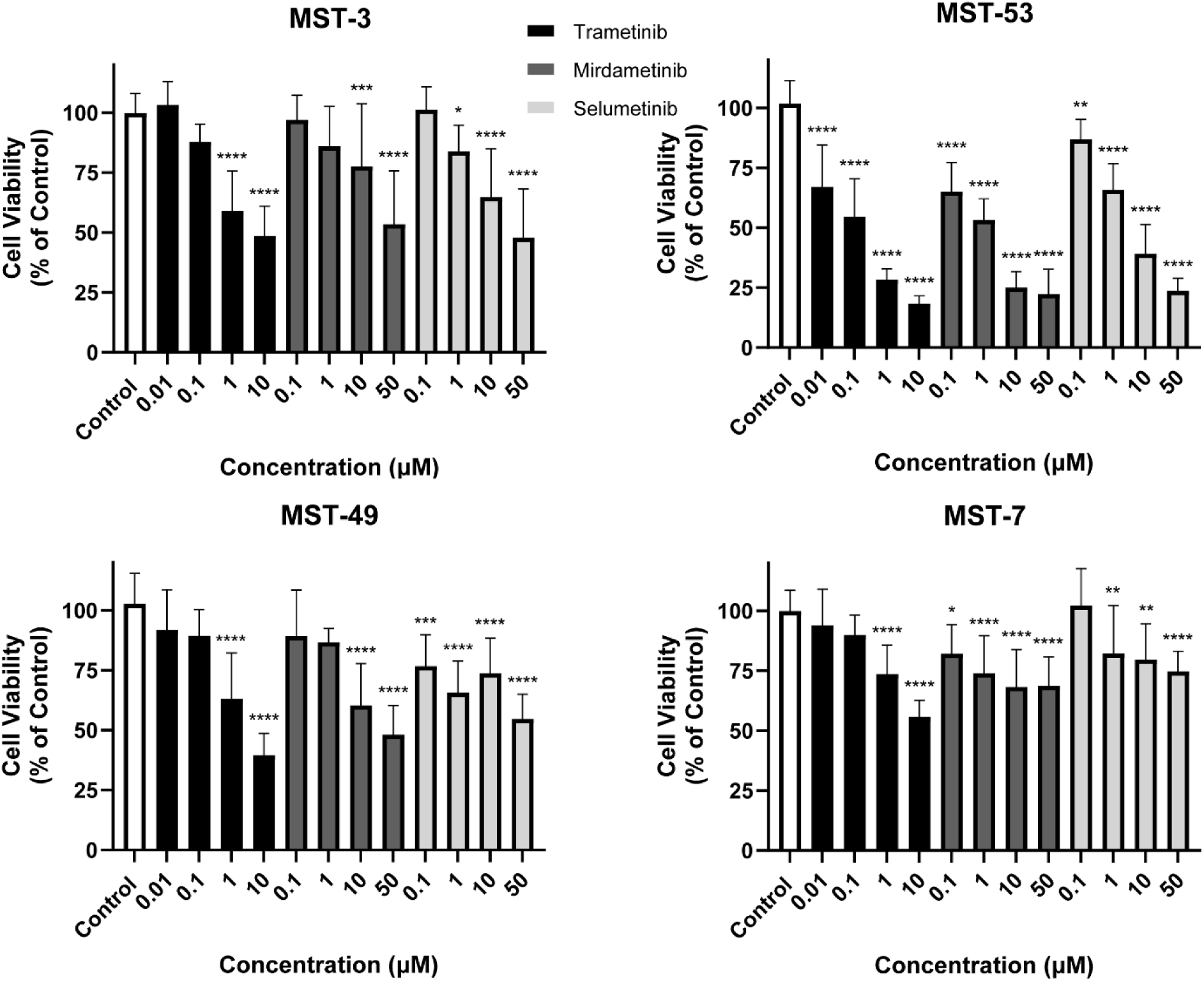
MEK small molecule blockade inhibits murine sinonasal tumor cell proliferation *in vitro*. Murine sinonasal tumor cell lines including ONB (MST-3), adenocarcinoma (MST-53), poorly differentiated carcinoma (MST-49), and SNSCC (MST-7) were treated with various doses of MEK inhibitors including Trametinib, Mirdametinib, and Selumetinib. Tumor cell proliferation was assessed *in vitro* and inhibition of cell proliferation was noted in each cell line with all three MEK inhibitors. Data are ± SEM of three independent experiments done in triplicates. *P* values are calculated by ordinary one-way ANOVA with Dunnett’s test. Significant *P* values are in comparison with the control group. * *P* < 0.05, ** *P* < 0.01, *** *P* < 0.001, **** *P <* 0.0001.

We then evaluated the efficacy of MEK inhibitors in human models of sinonasal malignancies. First, we utilized an available cell line of human SNSCC termed SCCNC1^16^. MEK inhibitors significantly inhibited SCCNC1 tumor cell proliferation (**Figure 6**, *P* < 0.05). Human models of sinonasal adenocarcinoma are sparse. We generated a robust human organoid model of sinonasal adenocarcinoma (termed NCI-089) from a patient who was found to have sinonasal intestinal-type adenocarcinoma with *TP53* and *NF1* tumor mutations (**Supplementary Figure 7**). MEK inhibitors significantly inhibited human NCI-089 organoid growth (**Figure 6**, *P* < 0.0001). Human models of ONB are also sparse. We generated two short-term human organoid models of olfactory neuroblastoma (termed NCI-546 and NCI-569) (**Supplementary Figure 8**). MEK inhibitors significantly inhibited early passage human ONB cell growth (**Figure 6**).

**Figure 6.**
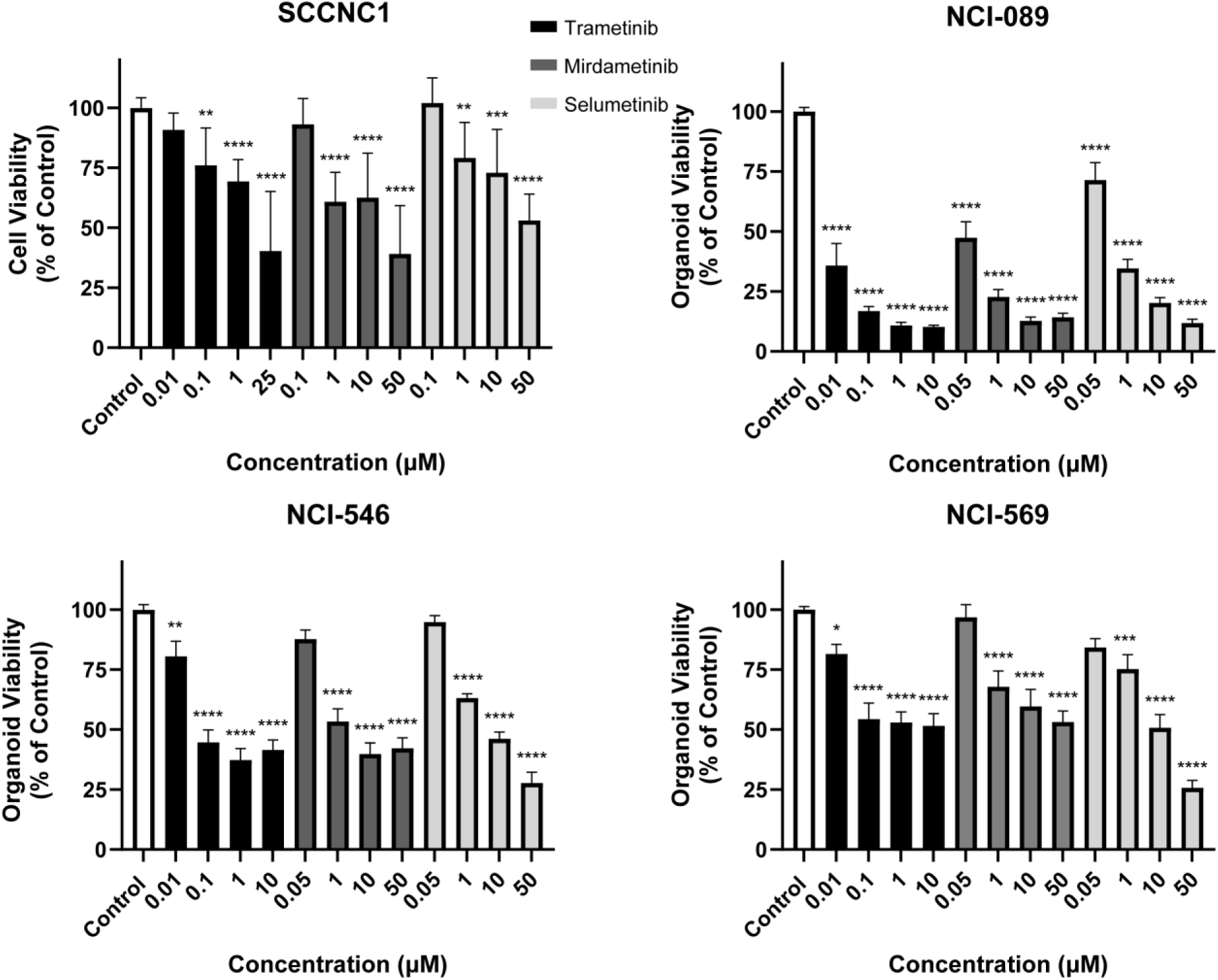
MEK small molecule blockade inhibits human sinonasal tumor cell proliferation *in vitro*. Human sinonasal tumor cell line SCCNC1, organoid models of sinonasal adenocarcinoma (NCI-089), and two ONB organoid models (ONB-546 and ONB-569) were treated with various doses of MEK inhibitors including Trametinib, Mirdametinib, and Selumetinib and tumor cell proliferation was assessed *in vitro*. Inhibition of cell proliferation was noted in each human model with all three MEK inhibitors. Data are ± SEM of three independent experiments done in triplicates. *P* values are calculated by ordinary one-way ANOVA with Dunnett’s test. Significant *P* values are in comparison with the control group. * *P* < 0.05, ** *P* < 0.01, *** *P* < 0.001, **** *P <* 0.0001.

## Discussion

Here, we performed the first sinonasal *in vivo* CRISPR screen in mice and surprisingly, rather than unique, distinct sgRNA mutational profiles across sinonasal tumor subtypes, common recurrent *NF1*/*RASA1*/*TP53* co-mutations across the major sinonasal tumor subtypes including SNSCC, ONB, adenocarcinoma, and SNUC were identified (**Figure 2-3**). We also developed many of the first reported mouse cell lines for these sinonasal tumor types and together with human cell line and newly developed human organoid models, elucidated the key function of downstream MEK signaling across sinonasal tumorigenesis. Collectively, these results suggest that loss of NF1 and RASA1-mediated Ras-GAP activity leads to Ras activation and downstream MEK signaling, which is a potential common target throughout the major sinonasal tumor subtypes (**Supplementary Figure 9**).

While *TP53* mutations are amongst the most common in sinonasal tumors^9–13^, *NF1* mutations have also been reported in multiple other sinonasal tumor types including SNSCC, sinonasal adenocarcinoma, and ONB^12,17–19^. Interestingly, a recent study of sinonasal adenocarcinoma whole genome sequencing reported that in addition to frequent *TP53* and *NF1* mutations, there were notable copy number variant (CNV) loss at 5q14.3-q15 where *RASA1* is located in 6/16 patients (38%)^12^. Additionally, *NF1* and *RASA1* co-mutations have been reported in a subset of non-small cell lung carcinomas^20^. Thus, human genomics studies are supportive that alterations in the molecular mechanism proposed in our study are observed and may be an important molecular target in human sinonasal malignancies. In addition to mutation and CNV analyses, it will be important in future studies to investigate other mechanisms such as DNA methylation and chromatin regulatory alterations that may also impact expression of these genes in sinonasal tumors. Indeed, loss of chromatin regulator proteins such as SMARCB1, SMARCA4, and ARID1A are emerging recurrent molecular mutations in human sinonasal tumors^2,21–24^. Our screen identified mutations across the entire SWI/SNF complex rather than only in SMARCB1, SMARCA4, and ARID1A, which suggests that in murine sinonasal tumors mutations across these SWI/SNF complex members may have similar downstream consequences.

Using *NF1/TP53^flox/flox^* mice, we found that loss of these two genes alone is sufficient to incite sinonasal tumor development. However, the fact that sinonasal tumors from *NF1/TP53^flox/flox^* mice were smaller and took ∼12 months to develop, rather than 3-5 months as seen in the AAV5-*TP53*-sgRNA screen, suggests that other mutations identified in the screen such as *RASA1* and DNA methylation and chromatin regulatory mechanisms may be important for accelerating sinonasal tumor growth. Compared to other genetically engineered mouse models (GEMM) in other organs driven by oncogenes, which may have a more rapid onset, the mouse models we developed here are incited by loss of tumor suppressor genes. Aside from isocitrate dehydrogenase-2 (IDH2) activating mutations in SNUC, activating mutations have not been regularly identified in sinonasal tumors. Thus, the relatively slower growth pattern of these tumors observed in the CRISPR screen and in *NF1/TP53^flox/flox^* mice driven by loss of tumor suppressor genes may resemble human sinonasal tumor growth patterns.

Here, we used small molecule MEK inhibitors including Trametinib and Selumetinib (both of which are FDA-approved) as well as Mirdametinib which is currently under investigation in clinical trials^25^. These drugs demonstrated efficacy over an array of murine and human sinonasal *in vitro* tumor models. Of these three inhibitors Trametinib was more effective at lower concentrations than Selumetinib and Mirdametinib. In addition to a direct anti-tumor effect, Trametinib has been reported to sensitize a variety of tumors including head and neck cancer to immunotherapy^26–28^. As the murine models in this study were generated in mice with a C57BL/6 background retaining an intact immune system, it will be interesting to investigate combinatorial Trametinib and immunotherapy approaches in these syngeneic mouse sinonasal tumor models in the future.

There are limitations to this study. While our approach discerned initial key inciting mutational events in sinonasal tumor development, our approach may not identify downstream mutations that may occur after these initial events. Secondly, MEK signaling is downstream of multiple signaling pathways. While small molecule inhibition of MEK may not be entirely specific for NF1/RASA1 downstream signaling, the broad efficacy across multiple murine and human *in vitro* models of various histologic subtypes also highlights the broad translational potential of MEK inhibition for sinonasal malignancies.

---

**Supplementary Figure 1.**
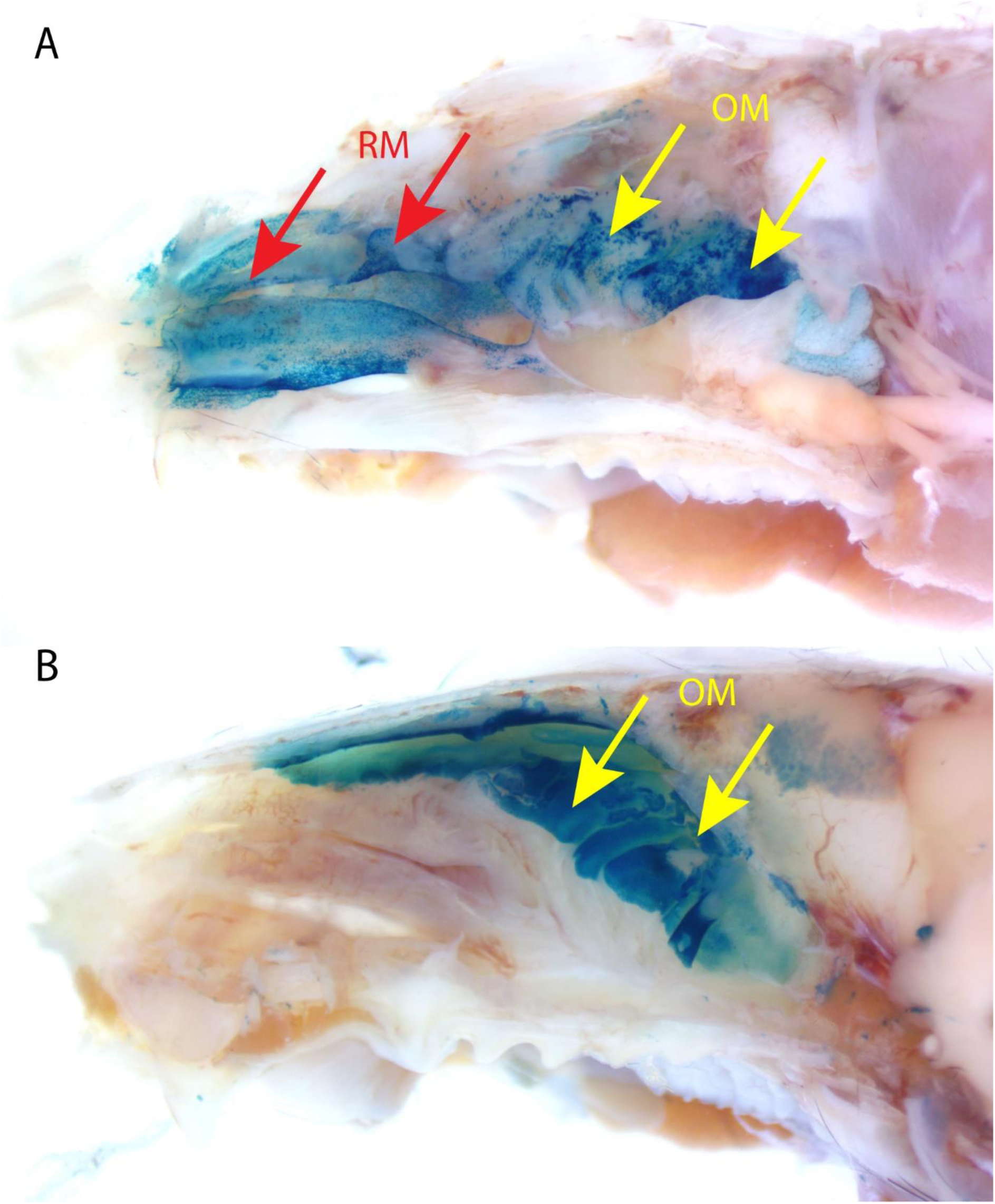
AAV5-Cre broadly transduces sinonasal cells across the olfactory and respiratory epithelium. **(A)** Sinonasal instillation of AAV5-Cre was performed in R26R mice. X-gal staining (blue) reveals representative transduction across the olfactory mucosa (OM, yellow arrows) and the respiratory mucosa (RM, red arrows). **(B)** For reference, OMP-Cre mice were crossed with R26R mice and X-gal staining demonstrates the location of the olfactory mucosa (yellow arrows).

**Supplementary Figure 2.**
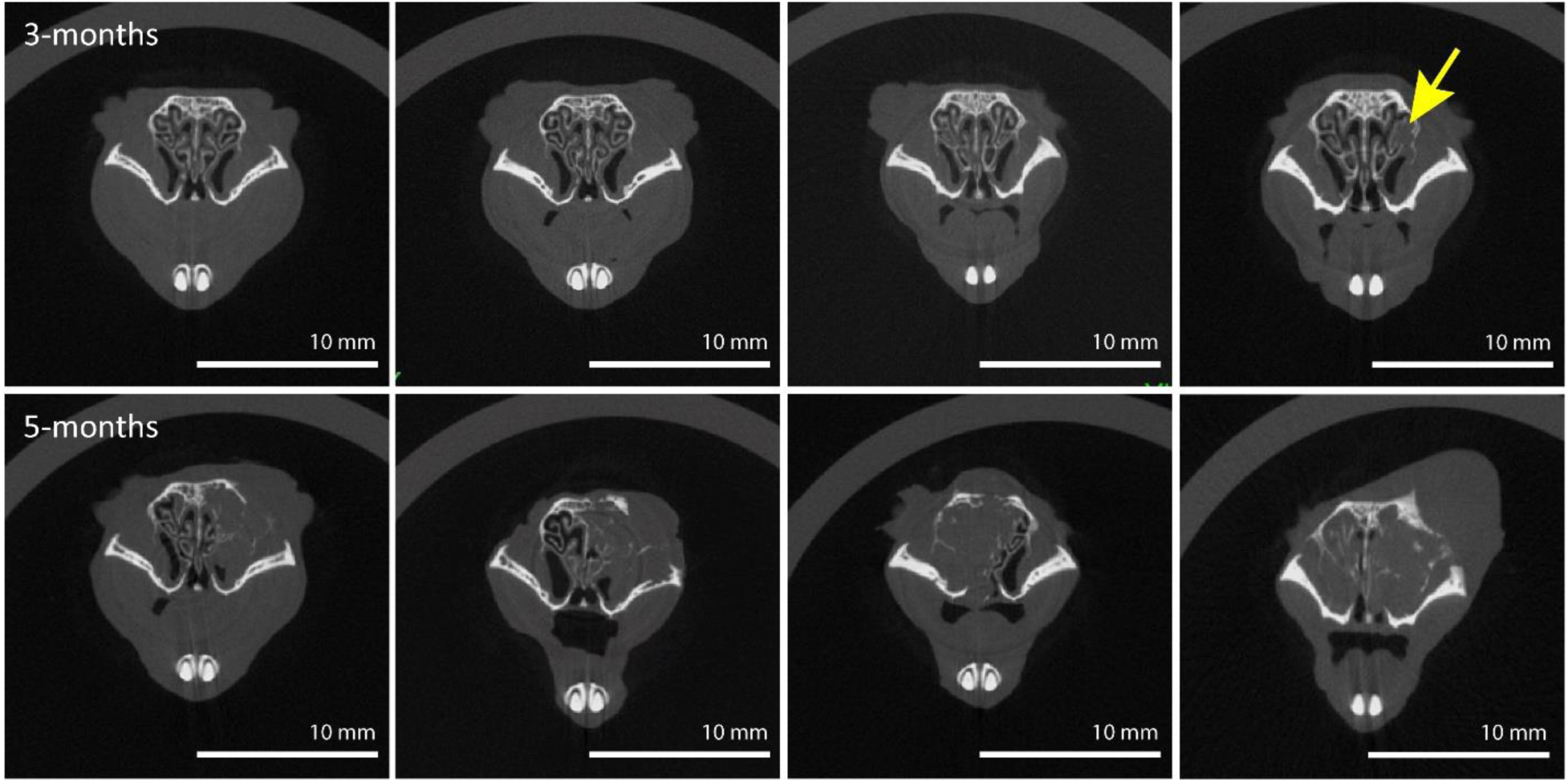
Sinonasal tumor formation after AAV5-*TP53*-sgRNA library instillation in H11*^Cas9^* mice on microCT. Thirty-three H11*^Cas9^* were exposed to the AAV5-*TP53*-sgRNA library. Initial signs of tumor formation were noted as early as 3 months post-instillation on microCT **(top row)** with large tumor formation noted around 5 months **(bottom row)**. Twenty-eight of these thirty-three H11*^Cas9^* mice developed a sinonasal tumor.

**Supplementary Figure 3.**
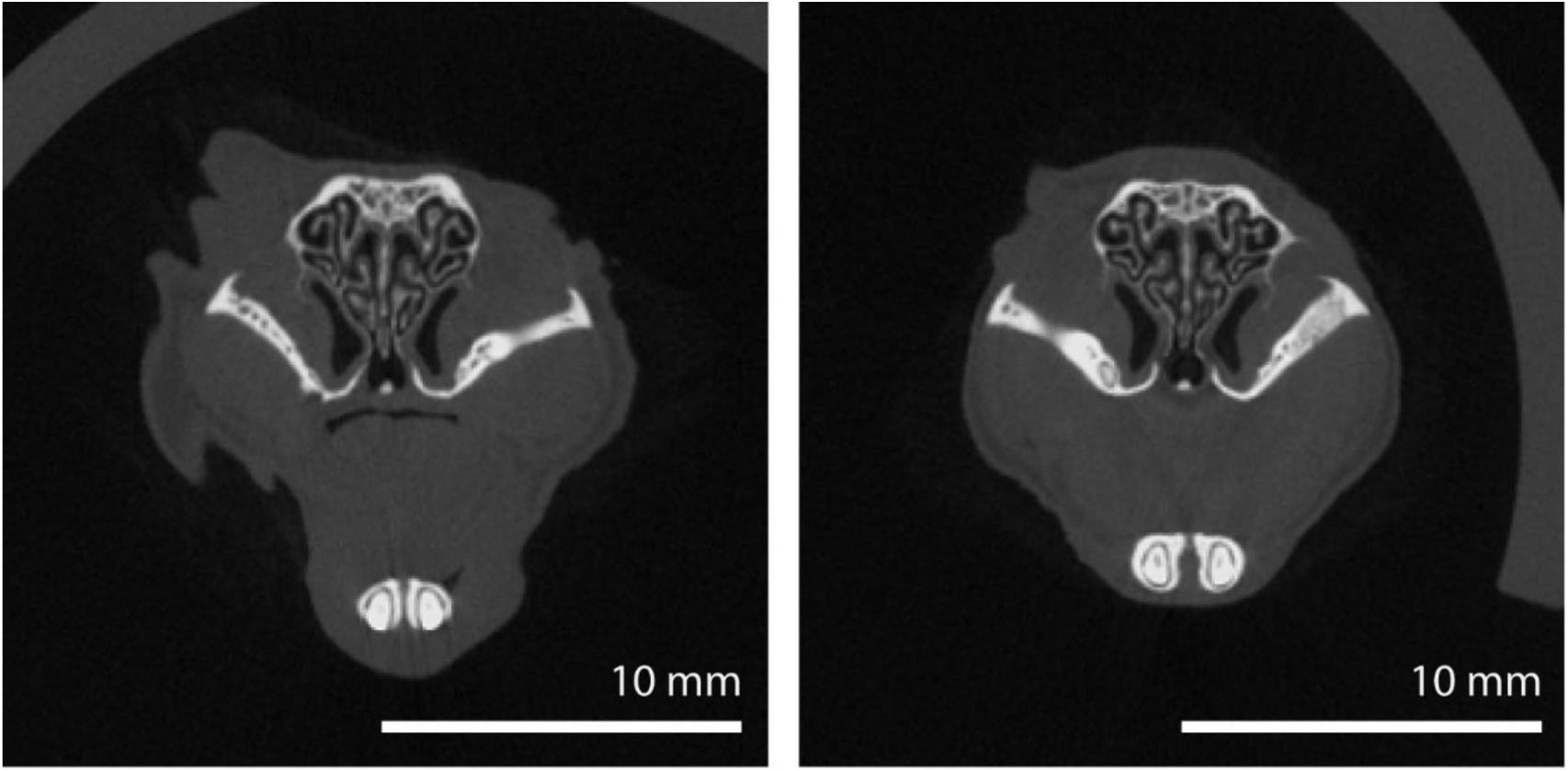
No sinonasal tumor formation observed after AAV5-*TP53*-null instillation in H11*^Cas9^* mice on microCT. Twenty-seven H11*^Cas9^* mice were exposed to AAV5-*TP53*-null control virus. No tumor formation was observed on microCT within 16 months after administration in any of these control mice. Representative coronal images from two mice are shown here.

**Supplementary Figure 4.**
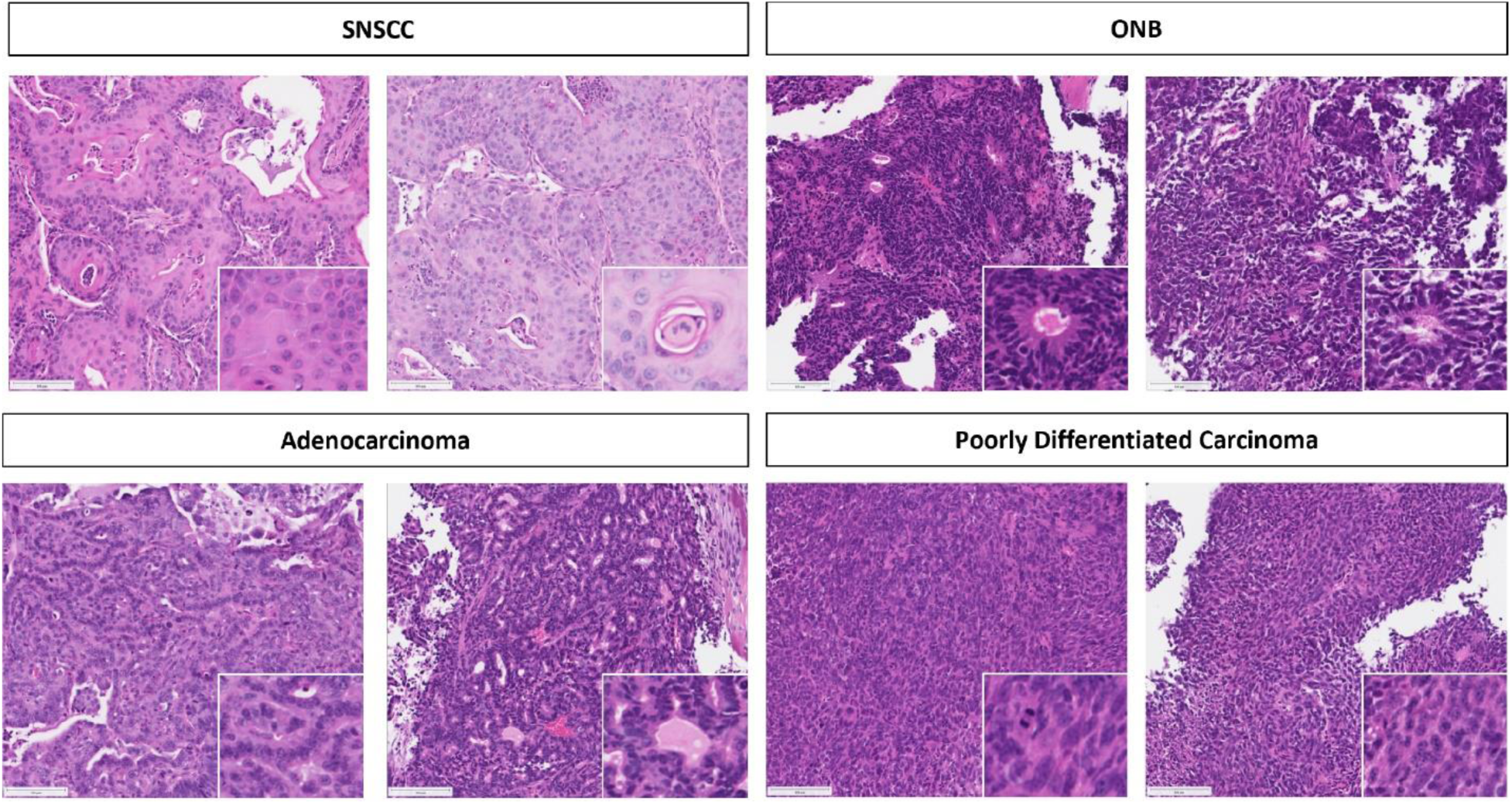
Histologic demonstration of murine sinonasal tumor types. Hematoxylin & Eosin staining demonstrated histologic findings consistent with SNSCC, ONB, adenocarcinoma, and poorly differentiated carcinoma. Representative images from two tumors from each tumor type are shown along with high magnification inset of key pathologic features. Scalebar equals 50µm. SNSCC – sinonasal squamous cell carcinoma, ONB – olfactory neuroblastoma.

**Supplementary Figure 5.**
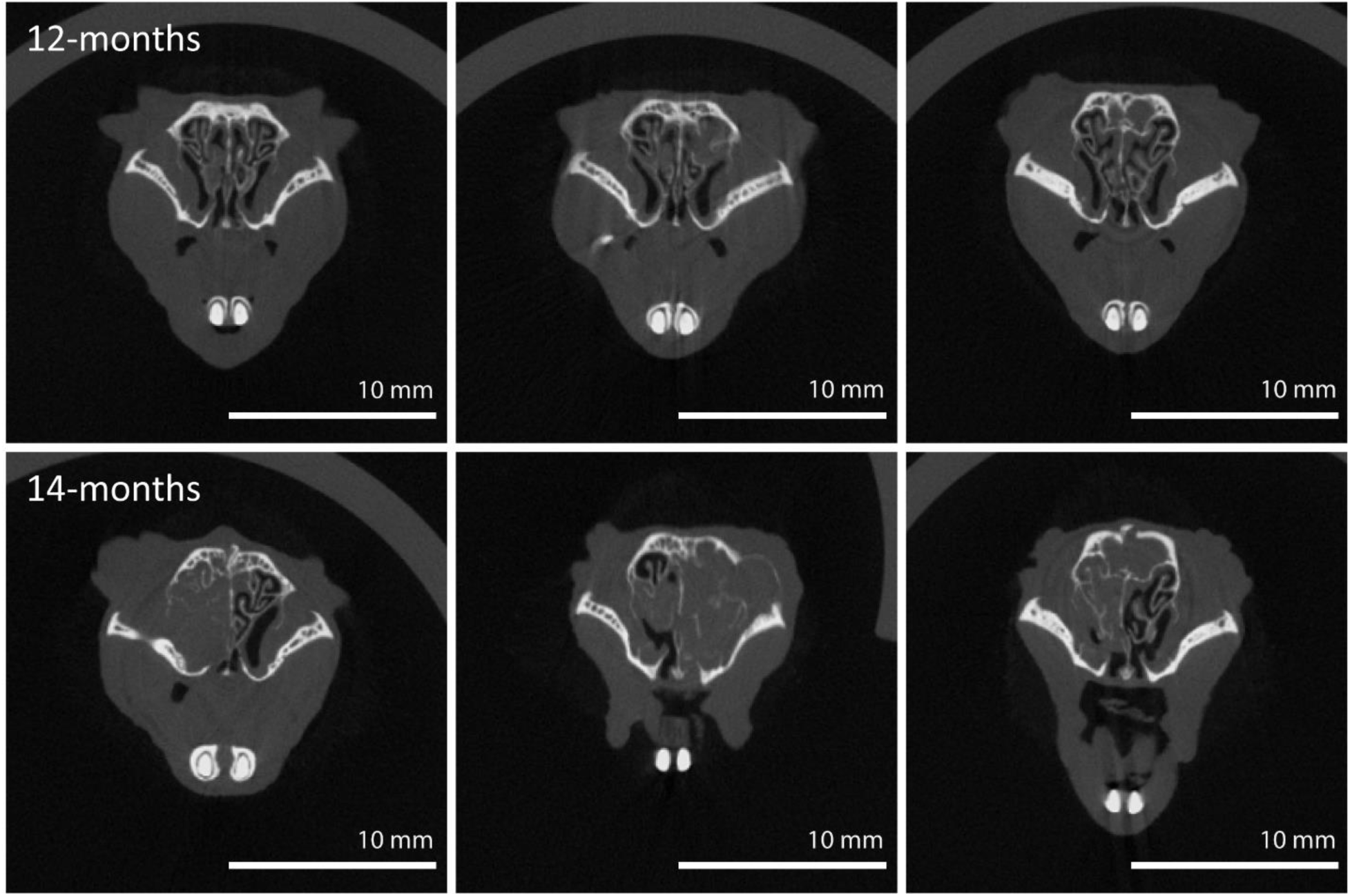
Sinonasal tumor formation after AAV5-Cre instillation in *NF1/TP53^flox/flox^* mice on microCT. Twenty *NF1/TP53^flox/flox^* mice were exposed to AAV5-Cre. Initial signs of tumor formation were noted around 12 months post-instillation on microCT **(top row)** with slow growth noted around 14 months **(bottom row)**. Thirteen of these twenty *NF1/TP53^flox/flox^*mice developed a sinonasal tumor.

**Supplementary Figure 6.**
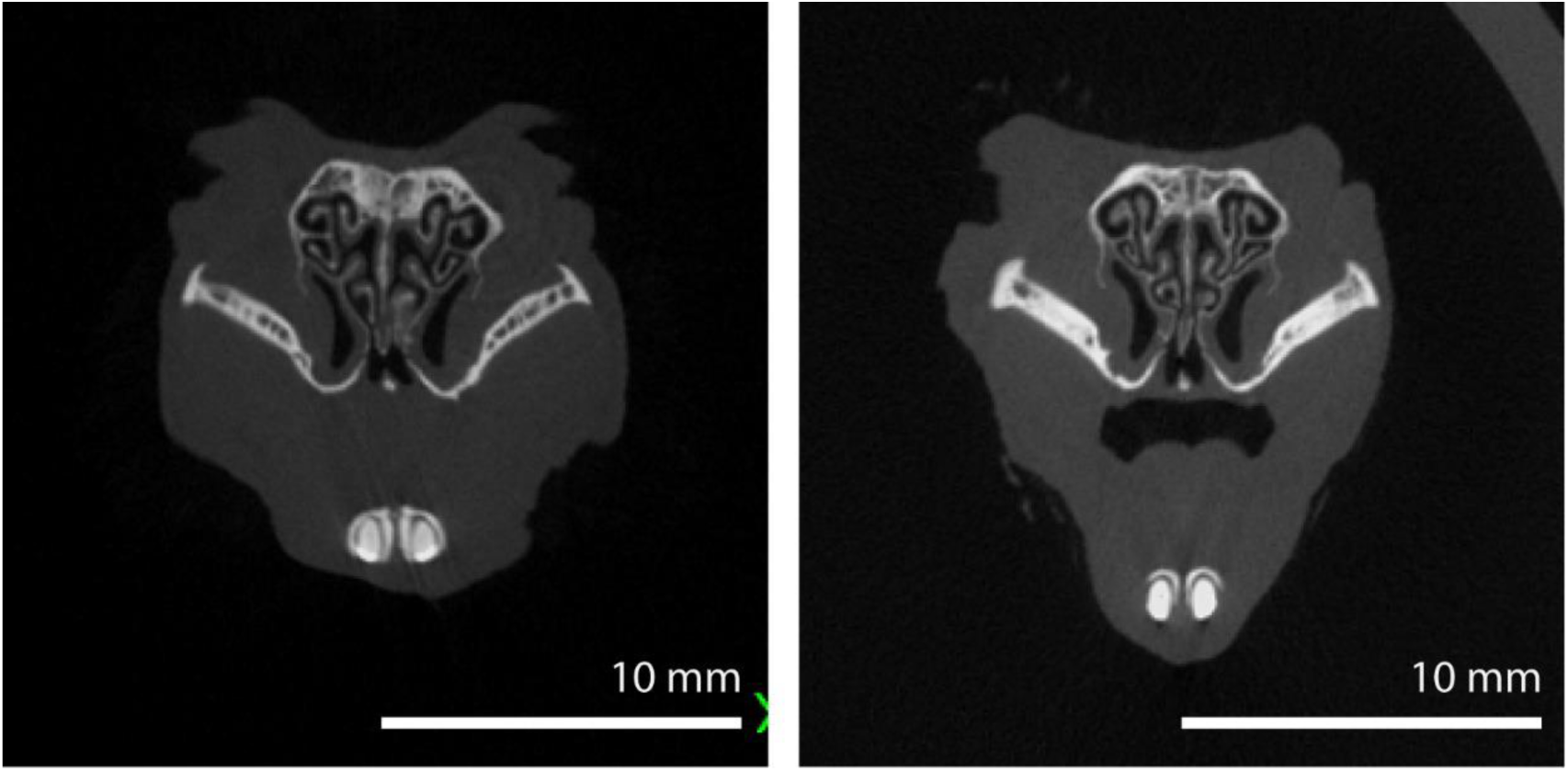
No sinonasal tumor formation observed after AAV5-null instillation in *NF1/TP53^flox/flox^* mice on microCT. Fifteen *NF1/TP53^flox/flox^* mice were exposed to AAV5-null control virus. No tumor formation was observed on microCT within 16 months after administration in any of these control mice. Representative coronal images from two mice are shown here.

**Supplementary Figure 7.**
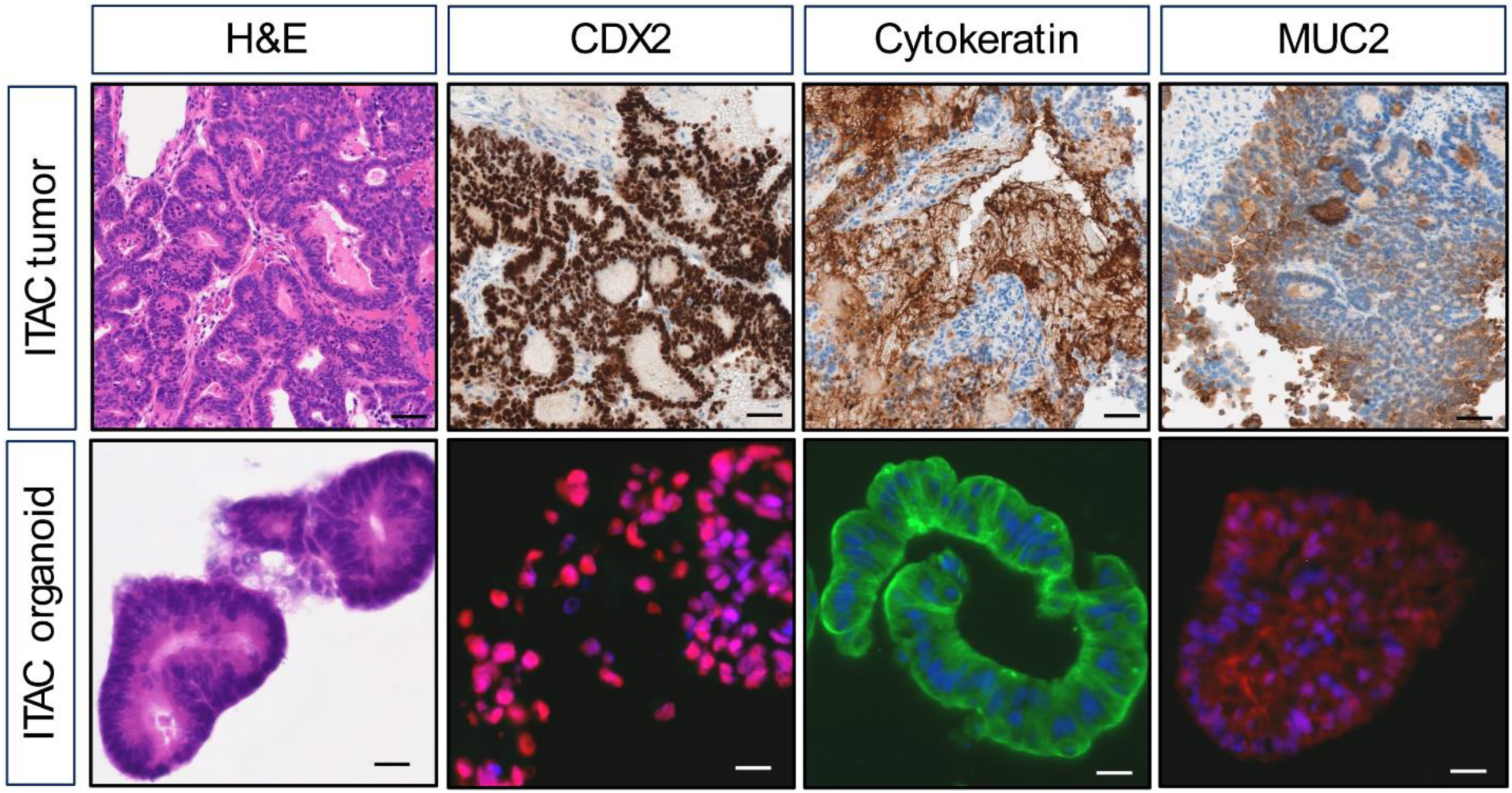
Histological characterization of human sinonasal intestinal type adenocarcinoma tumor tissue and organoids. (Top row) Human sinonasal intestinal type adenocarcinoma (ITAC) tumor tissue from which organoids were derived demonstrate histologic patterns and expression of markers of sinonasal ITAC including CDX2, cytokeratin, and MUC2. **(Bottom row)** Sinonasal ITAC organoids demonstrate histologic characteristics and expression of key markers including CDX2, cytokeratin, and MUC2. Scale bars for H&E and immunohistochemistry are 20 µM while scale bars in immunofluorescence are 200 µM.

**Supplementary Figure 8.**
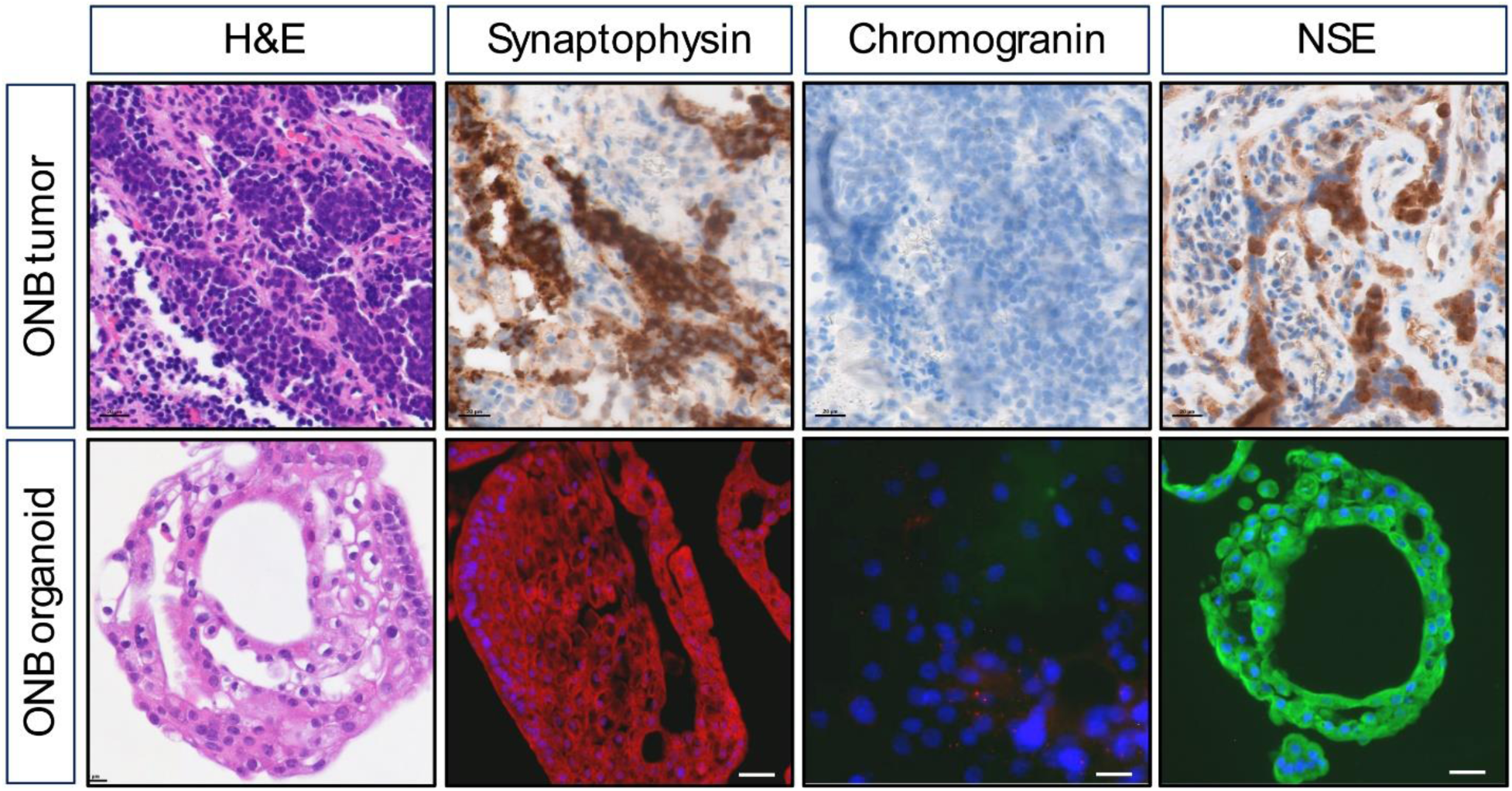
Histological characterization of human ONB tumor tissue and organoids. (Top row) Human olfactory neuroblastoma (ONB) tumor tissue from which organoids were derived demonstrate histologic patterns and expression of markers of ONB including synaptophysin and NSE. This particular tumor had low chromogranin expression. **(Bottom row)** ONB organoids demonstrate histologic characteristics and expression of key markers including synaptophysin and NSE with low chromogranin expression as seen in the primary tumor sample. Scale bars for H&E and immunohistochemistry are 20 µM while scale bars in immunofluorescence are 200 µM.

**Supplementary Figure 9.**
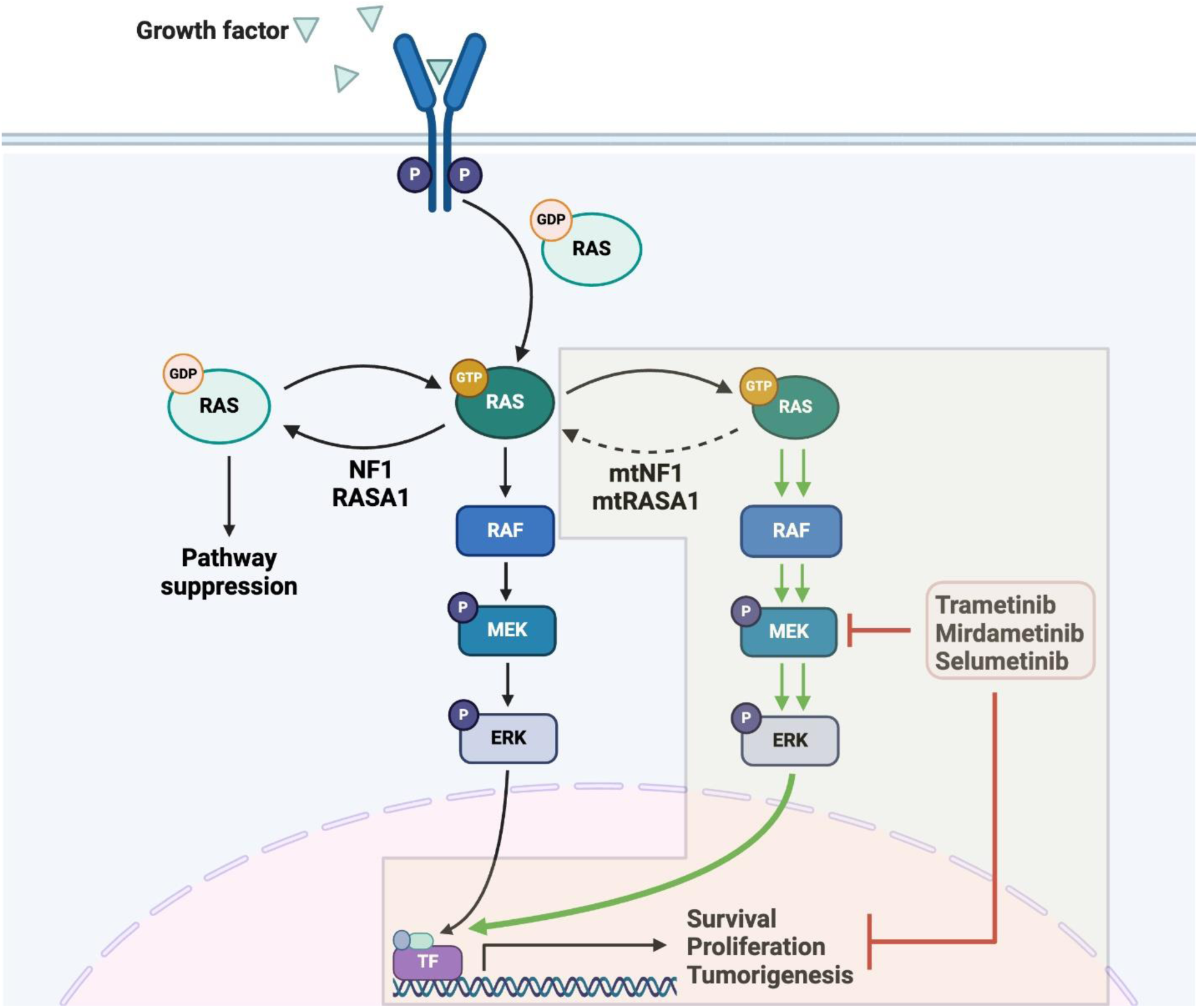
Loss of NF1 and RASA1-mediated Ras-GAP activity leads to Ras activation and downstream MEK signaling. Therefore, targeting MEK signaling with small molecule inhibitors such as Trametinib, Mirdametinib, and Selumetinib may be a potential common target in major sinonasal tumor subtypes.

**Supplementary Table 1.** List of sgRNA targets in the AAV5-*TP53*-sgRNA library.

**Supplementary Table 2.** Summary of genes with a lower frequency of sgRNA perturbation.

