## Supplementary material for "*In vivo* CRISPR screening identifies *NF1/RASA1/TP53* co-mutations and downstream MEK signaling as a common key mechanism of sinonasal tumorigenesis": Methods

### Materials and Methods

#### Animal Care and Use

National Cancer Institute (Bethesda, MD) guidelines for the care and use of laboratory animals were followed and all experiments were approved by the Animal Care and Use Committee at the National Institutes of Health.

#### Localization of Murine Sinonasal Cavity Viral Transduction

R26R mice were obtained from The Jackson Laboratory (strain #003474). Various serotypes of adeno-associated virus (AAV)-Cre vectors were obtained from Vector Biolabs (Malvern, PA) including AAV5-Cre (Cat #7088). R26R mice were briefly anesthetized with isoflurane and nasal instillation of 15 µL of viral-Cre once was performed daily for three days. Mice were euthanized 12 days later and dissected heads were fixed in 0.2% glutaraldehyde/PBS. The specimens were rinsed in detergent (0.02% Igepal, 0.01% Sodium Deoxycholate, and 2 mM MgCl2 in 0.1 M phosphate buffer [pH 7.5]) for 15 minutes, repeated 3 times. This was followed by incubation in X-gal staining solution (1 mg/mL X-gal [Invitrogen 15520034], 0.02% Igepal, 0.01% Sodium Deoxycholate, 5 mM Potassium Ferricyanide, 5 mM Potassium Ferrocyanide, and 2 mM MgCl2 diluted in 0.1 M phosphate buffer [pH 7.5]) for 1-2.5 hours. The heads were rinsed in PBS for 15 minutes 2 times and stored in 4% paraformaldehyde at 4°C. As a reference for the olfactory epithelium, R26R mice bred with olfactory marker protein (OMP)-Cre mice (JAX strain #006668) and underwent an identical X-gal staining procedure.

#### CRISPR Knockout Library Generation

To generate a CRISPR knockout library, 167 genes were selected from genes implicated in sinonasal and head and neck cancer tumorigenesis as well as prior *in vivo* CRISPR screening approaches in other organs (**Supplementary Table 1**)[1-3]. Up to five guide RNAs per gene were designed with four guides coming from the Brie library[4] and the fifth guide for each gene designed using sgRNA Scorer[5,6]. Ten non-targeting guides were also added. In total 835 guide RNAs were included in the library and a DNA oligonucleotide pool encoding these guide RNAs was obtained from Twist Biosciences. An AAV backbone carrying Cre-GFP, guide RNA cloning site, and Trp53 guide RNA was generated. Briefly, Addgene-61591 was digested with AgeI/KpnI and PCR fragments encoding Cre recombinase and GFP were assembled into this digested backbone. Subsequently, this plasmid was digested with NotI/KpnI and a synthesized guide RNA cloning cassette was assembled in this digested backbone. Finally, a cassette encoding the U6 promoter and a guide RNA targeting Trp53 (AGGAGCTCCTGACACTCGGA) was cloned into the previous vector by digesting that vector with XhoI/XbaI[7]. All cloning was performed using isothermal assembly[8]. The sgRNA library was cloned into the AAV backbone using HiFi Builder assembly (New England Biosciences) and sgRNA representation was assessed using high throughput Illumina sequencing. This was termed the AAV5-*TP53*-sgRNA library and the control which included the *TP53* sgRNA in the viral backbone was termed AAV5-*TP53*-null. Large-scale amplification and viral purification for *in vivo* use was performed by Vector Biolabs.

#### ***In Vivo*** **Sinonasal CRISPR Screen and Tumor Generation**

H11$^{Cas9}$ knock-in mice, which constitutively express Cas9 protein, were obtained from The Jackson Laboratory (strain #028239). Thirty-three 8-week-old H11$^{Cas9}$ mice (16 male and 17 female) underwent 3 days of nasal instillation of 15µL of AAV5-*TP53*-sgRNA library as described above. The viral titer was at least 3.6 x $10^{12}$ genome copies per mL (GC/mL) and lower viral titer dilutions uncommonly led to tumor development. For controls, 27 (14 male and 13 female) H$^{11}$ Cas9 mice were nasally instilled with AAV5-*TP53*-null as described above for 3 days. Twenty 8-

week-old (10 male and 10 female) *NF1/TP53*$^{flox/flox}$ mice (from Dr. Karlyne M. Reilly, NIH) underwent 3 days of nasal instillation of 15µL of AAV5-cre (Cat #7088, Vector Biolabs) with a viral titer of at least 5.0 x $10^{12}$ GC/mL. For controls, 15 (9 male and 6 female) *NF1/TP53*$^{flox/flox}$ mice underwent 3 days of nasal instillation of 15µL of AAV5-null (Cat #7028, Vector Biolabs). H11$^{Cas9}$ and *NF1/TP53*$^{flox/flox}$ mice initially underwent microCT imaging (Micro-CT Scanner Quantum GX) at 3 and 12 months, respectively. Mice were anesthetized with isoflurane, and images were acquired with 2 min scans at 36 mm FOV, 90 kV, with 88 mA of current. Mice were followed with serial microCT until study completion.

**Tumor histologic subtyping and immunohistochemistry**

Mice were euthanized once meeting experimental endpoints or by veterinary recommendation. Dissection was performed and tumor tissue first removed from the sinonasal cavity was snap frozen and later underwent DNA isolation for targeted sequencing. Patient tumor tissues were collected in saline from the operating room and a portion of it was snap frozen for DNA extraction. A small portion of murine and human tumor tissue was fixed in 10% neutral buffered formalin overnight and stored in 70% ethanol until processing. Fixed tumor samples were embedded in paraffin and 5µm sections were H&E stained (Histoserve Inc.). Immunohistochemistry was performed for synaptophysin (Abcam, ab32127), neuron-specific enolase (Santa Cruz, sc-21738), Chromogranin A (Abcam, ab254322), p63 (Abcam, ab124762), cytokeratin AE1/AE3 (SantaCruz,sc-81714), CDX-2 (Abcam, ab76541), and MUC-2 (Invitrogen, MA5-32654) as indicated on Leica bond automated stainer using 3’-‘3 diaminobenzidine tetrahydrochloride hydrate detection system (Bond Polymer Refine Detection, Leica Biosystems, DS9800). Slides were scanned at 40X or 20X magnification is with Vectra Polaris slide scanner. H&E, IHC slides, and digital images were blinded and evaluated by a veterinary pathologist (E.E.) at the Molecular

Histopathology Laboratory at the Frederick National Laboratory for Cancer Research and histopathology of each murine tumor determined.

**Targeted Sequencing**

Tumor DNA was isolated and concentration was measured by Qubit 1x dsDNA-HS assay kit (Invitrogen, cat # Q32231). 200 ng of DNA were sheared in 20 µL 1X Low TE buffer by Covaris Instrument (E220 PLUS) to ~ 175 bp. Shearing was done in 8 AFA-TUBE TPX Strip (Covaris #520292) with following parameters - duty factor 25%; peak incident power 220; cycle burst 50; time - 200 sec at 4ºC.

To generate a custom capture library for targeted genomic sequencing, genomic coordinates for each guide RNA were obtained using the BLAT tool against the Mm10 mouse genome in the UCSC Genome Browser. Approximately 100 nucleotides on each side of the target site was used as the basis of the probe for custom capture and the library was prepared (Agilent Technologies, Santa Clara, CA). The Agilent SureSelect XT HS Target Enrichment System for Illumina paired-end sequencing library protocol was used for library preparation (Agilent Technologies, Santa Clara, CA). Agilent custom mouse capture kit # 5191-6900 (Design/ ELID #3296641) was used. DNA Lo Bind Tubes, 1.5-mL PCR clean (Eppendorf #022431021) or 96 well plates were used to process the samples.

Agilent SureSelectXT-HS library prep kit (Agilent # G9702A) was used to prepare the library for each sheared DNA sample. DNA fragment ends were repaired and adenylated at the 3' end and then indexed with individual Index and adaptor ligated. These libraries were then amplified (pre-capture PCR amplification - 98°C 2 minutes, 8 cycles – 98°C, 30 seconds 60°C, 30 seconds 72°C 1 minute then 72°C 5 minutes, 4°C hold) by Herculase II fusion enzyme provided with the Agilent library preparation kit. Adaptor ligated amplified libraries were purified by Ampure XP beads (Beckmann Coulter Genomics # A63882).

Samples were analyzed by Tapestation DNA-1000 screen tape (Agilent # 5067-5582) to check the size of the libraries (Agilent 4200 TapeStation # G2991 AA). Concentration of each library was determined by integrating area 200-500 bp. Then each library (~500 - 1000 ng) was hybridized with biotinylated custom mouse capture RNA baits (catalog # 5191-6900; Design/ELID # 3296641) in the presence of blocking oligonucleotides. Hybridization was done as follows - 95°C, 5 minutes 65°C, 10 minutes 65°C, 1 minute 60 cycles, 65°C 1 minute, 37°C 3 sec. then 65°C hold. Custom mouse capture baits were added to the libraries after pausing the thermal cycler after 5 minutes at 65ºC incubation. Capture and post capture steps were done by automation on Agilent-BRAVO Liquid handler (Model: G5563A). Bait-target hybrids were captured by streptavidin-coated magnetic beads (Dynabeads MyOne Streptavidin T1, Life Technologies, # 6560) for 30 minutes at room temperature. Then after a series of washes to remove the non-specifically bound DNA, captured library was eluted in nuclease free water and amplified (Post-capture PCR amplification - 98°C 2 minutes, 12 cycles– 98°C 30 seconds, 60°C 30 seconds, 72°C 1 minute, then 72°C 5 minutes, 4°C hold). Tapestation High sensitivity DNA screen Tape (Agilent # 5067-5587) was used to validate the size of the libraries.

Libraries were pooled together based on the TapeStation concentrations and dilute to 4nM with water. 5ul 0.2N NaOH was added to 5ul of pooled library to denature 5min at room temperature. 990ul HT1 buffer was added to make the denatured library concentration to 20pM. Then 200ul denatured library, 396ul HT1 buffer and 4ul denatured Phix control were mixed together. The final library concentration was about 6.7pM. 600ul of the final library was added to the MiSeq reagent cartridge and run on MiSeq 154 + 8 + 154 cycles. For each tumor sample, single nucleotide variants (SNVs) and INDELs were identified using Freebayes and mm10/GRCm38 genome assembly.

**Bioinformatics analysis**

Raw sequencing reads were processed with the nf-core/sarek pipeline v3.4.0[9] in tumor-only mode, using the mm10/GRCh38 reference genome. Sequence reads were aligned to the reference genome using BWA-MEM2, followed by GATK4 best practices to mark duplicates and base recalibration[10]. Somatic single nucleotide variants (SNVs) and small insertions and deletions (INDELs) were called using Freebayes[11] with default parameters. Variants detected by Freebayes were extracted with vcflib[11] to retain only INDELs, thus restricting the analysis to those mutational events that can be considered direct consequences of CRISPR/Cas9 editing. Variants effects were predicted using VEP (v110)[12]. Annotated VCF files were converted to MAF using vcf2maf[13]. Downstream data analysis and visualization were performed with maftools[14]. For Freebayes, only non-synonymous variants (i.e variants with predicted high/moderate consequence) were considered.

**Murine sinonasal tumor cell line generation**

Tumor samples were dissociated into a single cell suspension using the Miltenyi Biotec Tumor Dissociation Kit (130-096-730). Cells were plated onto gelatin coated (Stem Cell Technologies 07903) 6-well plates. Cells were cultured in a media consisting of Advanced DMEM/F12, Pen/Strep, GlutaMax (Thermofisher 12634010, 15070063, 35050061), 5% fetal bovine serum (Innovative Research, IGFBSERHI), 0.4 μg/mL hydrocortisone, 24 μg/mL adenine (Milipore Sigma H0888 and A2786), 10 ng/mL human recombinant epidermal growth factor, and 10 μM Y-27632 dihydrochloride (Stem Cell Technologies 78006 and 72304).

**Human sinonasal organoids generation**

Human organoids were established as previously (Zamuner et al., 2024)[15]. In brief, tumor tissues were dissociated into single cell suspensions using tumor dissociation kit per manufacturer instructions (Miltney biotech., 130-095-929). The tumor single cells were suspended in ice cold 70% basement membrane extract (BME, Bio-techne, 3533-010-02) and cultured as domes in 24 well plates. The patient derived organoid growth media for sinonasal adenocarcinoma organoids consisted of 50% L-WRN conditioned media (prepared from L-WRN expressing cell line, ATCC, CRL-3276) 50% advanced DMEM/F12 supplemented with 1X-B27, 1X pencillin streptomycin, 1X HPES, 1X glutamax (Thermofisher; 12634-010, 17504-001, 15140-122, 15630080, 35050-061), 1.25mM N-acetyl-L-Cysteine, 10mM Nicotinamide (Sigma; 72340, A9166), 0.5uM A83-01, 1uM SB202190 (SelleckChem; S7692, S1077) and 50 ng/ml hEGF (Peprotech, AF-100-15). For culturing organoids from olfactory neuroblastoma, media consisted additional components including 1uM prostaglandin E2, 1uM Forskolin, 0.3uM CHIR99021(SelleckChem; S3003, S1263, S2449), 10ng/ml FGF10 and 5ng/ml FGF2 (Peprotech; 100-26,100-18B). After two rounds of passages, ONB organoids were cultured as 2.5D lines on gelatin coated plates.

**Organoid immunofluorescence**

The established sinonasal adenocarcinoma organoid was cultured for several passages and revives after cryopreservation, whereas ONB organoids survived for 5 to 6 passages. Cultured organoids were harvested and fixed in 4% paraformaldehyde overnight at 4ºC. The fixed organoid pellet was processed for frozen sections and were stained for H&E (Histoserve Inc.,). Immunoflourescence staining of frozen sections was carried out with antibodies listed in the immunohistochemistry methods section using Alexa 594 and Alexa 488 secondary antibodies. (Life technologies; A-

11012, A-11001) and images were captured with EVOS cell imaging system (Thermofisher scientific).

**Cell viability experiments**

Murine cell lines were selected based on pathology type and treated with the MEK inhibitors Selumetinib, Mirdametinib, and Trametininb (Selleckchem S1008, S1036, S2673) at varying concentrations from 0.01 µM to 100 µM. SCCNC1 cells were kindly provided by Dr. Mario Hermsen[16]. Cells were plated into gelatin coated 96 well plates and the drugs administered 24 hours after. 72 hours after drug administration an XTT assay was done (R&D Systems 4891-025-K). Controls were treated with equivalent amounts of DMSO only.

For ITAC, drug sensitivity experiments were carried out as previously described[17]. In brief, two days before treatment with drugs, organoids were harvested and passaged into single cells and plated in 70% BME and cultured in growth media. Two days later, organoids were collected by digesting BME with dispase II (Sigma, D4693), filtered through 70uM nylon cell strainer and were counted. Organoids were resuspended in 5% BME with ice cold growth media 40,000 cells/well/100ul and seeded into 96 wells plate. Two hours later, different doses of MEK inhibitors were added to the organoids and 120 hours after the treating with drugs, organoid viability was assessed by measuring ATP levels using CellTiter-Glo3-D reagent (Promega, G9681).

ONB cell viabilities were performed as previously described[15]. About 10,000 single cells were seeded into 96 well plates and different doses of MEK inhibitors were added after overnight attachment. Cell viability was assessed 72 hours after treating with drugs using CellTiter-Glo 2.0 (Promega, G9241) following manufacturer’s instructions. DMSO was used as vehicle control and data was normalized to vehicle (100% viability) and baseline control (0%) 0.1% TritonX-100. Kill curves were produced by fitting the lines with log inhibitor versus normalized response-variable slope using GraphPad Prism software.
